# Factors Affecting Protein Recovery During Hsp40 Affinity Profiling

**DOI:** 10.1101/2022.03.31.485989

**Authors:** Maureen R. Montoya, Guy M. Quanrud, Liangyong Mei, José L. Moñtano, Caleb Hong, Joseph C. Genereux

## Abstract

Hsp40s, also termed J-domain proteins, play a central role in cellular protein homeostasis by promiscuously surveying the proteome for misfolded proteins. We have exploited this property to develop Hsp40 affinity profiling as a method for identifying proteins that misfold in response to cellular stresses. In this assay, we use the Hsp40 ^Flag^DNAJB8^H31Q^ as our recognition element for misfolded proteins. This protein is exogenously introduced into cells, promoting interactions without regard for native protein clients. Herein, we evaluate potential approaches to improve the performance of this assay. We find that although intracellular crosslinking increases recovery of protein interactors, this is not enough to overcome the relative drop in DNAJB8 recovery. While the J-domain promotes Hsp70 association, it does not affect the yield of protein association with DNJAB8 under *basal* conditions. By contrast, crosslinking and J-domain ablation both substantially increase relative protein interactor recovery with the structurally distinct Class B Hps40 DNAJB1 but are completely compensated by poorer yield of DNAJB1 itself. Cellular thermal stress promotes increased affinity between DNAJB8^H31Q^ and interacting proteins, as expected for interactions driven by recognition of misfolded proteins. DNAJB8^WT^ does not demonstrate such a property, suggesting that under stress misfolded proteins are handed off to Hsp70. Hence, we find that DNAJB8^H31Q^ is still our most effective recognition element for the recovery of destabilized client proteins following cellular stress.

Raw data is accessible through the PRIDE Archive at PXD030633.

## INTRODUCTION

Cellular health requires the maintenance of protein homeostasis to prevent the accumulation of misfolded proteins and proteotoxic species[1]. A variety of exposure agents can damage proteins, threatening this maintenance of proteome integrity[2]. To evaluate these threats, we recently introduced Hsp40 Affinity Profiling as a technology for identifying proteins that are destabilized by cellular toxicant exposure [3–5]. It exploits the capacity of Hsp40 family proteins (also called J-domain proteins) to recognize and bind to misfolded proteins [6,7]. In this assay, a human Hsp40 DNAJB8 is expressed in cells prior to challenging the cells with the toxicant of interest. Misfolded proteins accumulate on DNAJB8, so that after lysis, they can be co-purified and identified and quantified by quantitative mass spectrometry (**Figure 1**). A J-domain mutation, H31Q, was introduced into DNAJB8 to prevent handoff of misfolded proteins to Hsp70 and thus ensure accumulation on DNAJB8. The strong affinity for DNJAB8^H31Q^ for misfolded proteins allows stringent washing with RIPA buffer to remove non-specific interactors. This approach has enabled us to identify proteins that are sensitive to metals and to electrophilic herbicides.

**Figure 1:**
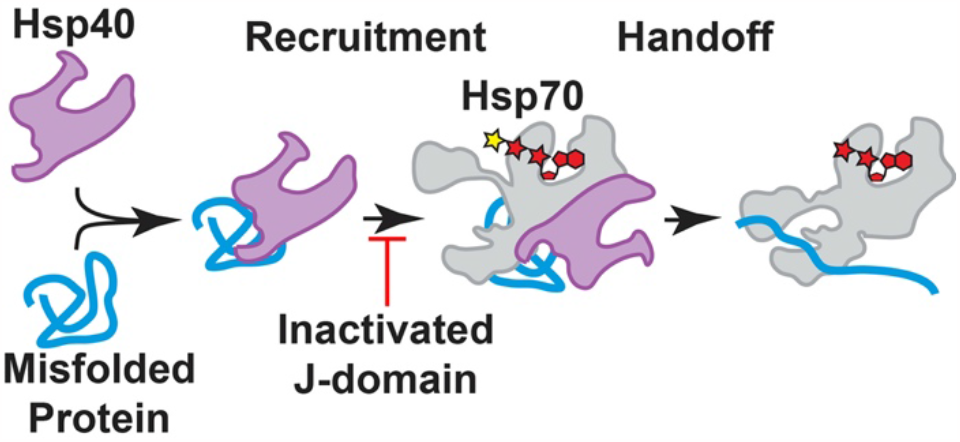
Hsp40 proteins recruit misfolded proteins to Hsp70 in a manned dependent on their J-domain.

Although our approach has been validated by its success, we have not established answers to key questions. Firstly, it is not clear that DNAJB8 is the only Hsp40 that could be used in this assay. Other Hsp40s might be able to be used to extract and assess the same misfolded proteins from biological samples, or perhaps even to target a complementary proteome. Secondly, although the H31Q mutation in the J-domain was introduced to avoid handoff of misfolded proteins to Hsp70, we do not know if it is necessary. Finally, we have not fully evaluated whether crosslinking in the cell provided improved misfolded protein recovery. Herein we consider these questions, finding that DNAJB8 requires neither crosslinking nor mutation to pull down its distinct proteome. However, under heat stress, the H31Q is necessary to observe increased Hsp40 affinity.

## MATERIALS AND METHODS

### Reagents

Biochemical reagents and buffer components were purchased from Fisher Scientific, VWR, or Millipore Sigma. Millipore water and sterilized consumables were used for all biochemical experiments.

### Molecular Cloning

DNAJB1 was amplified from cDNA derived from HEK293T cells (ATCC) using TRIzol (Thermo Fisher Scientific) and inserted into the pFlag.CMV2 vector by PIPE cloning[8] using Q5 polymerase and the primers:

5’ - CAGATCTATCGATGAATTCGCTATTGGAAGAACCTGCTCAAG -3’,

5’ - CTTGAGCAGGTTCTTCCAATAGCGAATTCATCGATAGATCTG - 3’,

5’ - GTAGTAGTCTTTACCCATGACCTTGTCGTCATCGTCTTTG - 3’, and

5’- CAAAGACGATGACGACAAGGTCATGGGTAAAGACTACTAC - 3’. The H32Q mutation was introduced into DNAJB1 using site-directed mutagenesis with the oligonucleotides

5’-CTACCAACCGGACAAGAACAAGGAGCCCGG-3’ and 5’-CCGGTTGGTAGCGCAGCGCC-3’.

^Flag^DNAJB8^WT^.CMV2, and ^Flag^DNAJB8^H31Q^.CMV2, and EGFP.pDest30 have been reported[9,10]. Constructs were analytically digested and sequenced (Retrogen) to confirm identity. All cloning enzymes and buffers were purchased from New England Biolabs and primers were purchased from IDT.

### Human Tissue Culture

These experiments were performed in HEK293T, which do not represent any specific human tissue type. However, proteostasis mechanisms tend to be highly conserved across euploidal cell lines. Therefore, it is likely that the general observations made here will hold, although specific clients might be handled differently. HEK293T cells were cultured in DMEM (Corning) supplemented with 10% fetal bovine serum (FBS; Seradigm), 2 mM L-Glutamine (Corning), and penicillin-streptomycin (100 IU/mL,100 µg/mL; Corning). Cells were transfected with plasmid DNA by the calcium phosphate method. Every experiment involving DNAJB8 used one 10 cm plate per condition. 4-plex experiments involving DNAJB1 used two 10 cm plates per condition to account for its lower overexpression. Where heat shock was applied, cells were placed in an incubator at the indicated temperature for 30 min. and then immediately harvested, washed, and the pellets frozen at –80 °C.

### Immunoprecipitation

Cells were harvested from confluent dishes at 36 h to 48 h post-transfection. If crosslinking was used, cells were incubated in the indicated concentration of freshly prepared dithiobis succinimidyl propionate (DSP) in 1% DMSO/PBS for 30 min with rotation at ambient temperature, and then quenched by addition of Tris pH 8.0 (to 90 mM final concentration) and rotation for 15 min. After crosslinking, or directly after harvest for experiments without crosslinking, cells were lysed for 30 min on ice in lysis buffer supplemented with fresh 1 x protease inhibitor cocktail (Roche). Unless otherwise indicated, lysis was performed in RIPA buffer (150 mM NaCl, 50 mM Tris pH 7.5, 1% Triton X100, 0.5% sodium deoxycholate, 0.1 % SDS). For low stringency experiments using DNAJB1, lysis was performed with 0.1% Triton x-100 in TBS (10 mM Tris pH 7.5, 150 mM NaCl). Lysate was separated from cell debris by centrifugation at 21,100 x g for 15 min at 4 °C. Protein was quantified by Bradford assay (Bio-Rad). Lysates were pre-cleared with 15 µL sepharose-4B beads (Millipore Sigma) for 30 min at 4 °C, followed by immunoprecipitation with 15 µL M2 anti-FLAG Magnetic Beads (Millipore Sigma) and overnight rotation at 4°C. Beads were washed four times with lysis buffer the next day for DNAJB8 or three days later for DNAJB1. Proteins were eluted from the beads by boiling in 30 µL of Laemmli concentrate (120 mM Tris pH 6.8, 60% glycerol, 12% SDS, brilliant phenol blue to color). About 17% of each eluate was reserved for silver stain analysis, and the remainder prepared for mass spectrometry.

### Silver Stain

Eluates were boiled for 5 min at 100 °C with 0.17 M DTT, loaded into 1.0 mm, 12% polyacrylamide gels, and separated by SDS-PAGE. Gels were rinsed in Millipore water for 5 min. Gels were left overnight in fixing solution (10% acetic acid, 30% ethanol), washed 3 x 20 min in 35% ethanol, sensitized (0.02% sodium thiosulfate) for 2 min, washed with Millipore water 3 x 2 min, and stained for 30 min to overnight in Ag staining solution (0.2% AgNO3, 0.076% formalin). Gels were washed 2 x 1 min with Millipore water and developed (6% sodium carbonate, 0.05% formalin, 0.0004% sodium thiosulfate) until bands reached desired intensity and imaged on a white-light transilluminator (UVP).

### TMT-MuDPIT

Immunoprecipitates were prepared for TMT-AP-MS according to standard protocols[11–13]. After TMT labeling, each TMT reaction was quenched with 0.4% ammonium bicarbonate. Labeled digests were combined and fractionated by SCX in line with a reversed-phase analytical column to enable two-dimensional separation prior to electrospray ionization. Peptides were analyzed using a LTQ Orbitrap Velos Pro in data-dependent mode. The top ten peaks from each full precursor scan were fragmented by HCD (stepped collision energies of 36%, 42%, 48% with 100 ms activation time) to acquire fragmentation spectra with 7500 resolving power at 400 m/z. Dynamic exclusion parameters were 1 repeat count, 30 ms repeat duration, 500 exclusion list size, 120 s exclusion duration, and 2.00 Da exclusion width. Peptide-spectra matches were evaluated by FragPipe[14] against the Uniprot human proteome database (Jun 11, 2021 release, longest entry for each protein) with 20429 sequences (including common contaminants) plus a full reversed sequence decoy set. Cysteine alkylation (+57.02146 Da) and TMT modification (+229.1629 on lysine and N-termini) were set as fixed modifications. Half tryptic and fully tryptic peptides were allowed, as were 2 missed cleavages per peptide. A mass tolerance of 1 Da for precursors and 20 ppm for fragments was allowed. Decoy proteins, non-human contaminants, immunoglobulins, and keratins were filtered from the final protein list.

### Gene Ontology

Selective interactors were analyzed by Panther 17.0 by comparison to all Homo sapiens genes by biological process. Ontologies were evaluated by False Discovery Rate based on Fisher’s Exact Test.

### Statistical Methods

TMT intensity ratios were analyzed using Excel. Box and whisker plots are presented with lines marking median values, X marking average values, boxes from the first to third quartiles, whiskers extending to minimum and maximum values (excluding outliers), and outliers defined at points greater than 1.5-fold the interquartile range beyond the first and third quartiles. Violin plots were generated in R using the ggplot2 library. For bait vs. mock experiments, Pearson’s R-derived t-statistics were used for determination of p-values[9]. q-values (qBH) were determined from p-values using Storey’s modification of Benjamini-Hochberg’s methodology[15,16], and adjusted to maintain monotonicity. For heat shock experiments with DNAJB8^H31Q^ and DNAJB8^WT^, integrated TMT reporter ion intensities of identified proteins were normalized to bait intensities.

## RESULTS

^*Flag*^*DNAJB8*^*WT*^ *specifically enriches hundreds of proteins in TMT-AP-MS*.

We originally incorporated the J-domain H31Q mutation into DNAJB8 to prevent hand-off of mutant proteins to Hsp70 (**Figure 1**), but we did not evaluate whether this mutation was necessary for the observed strong protein binding. To evaluate whether DNAJB8^WT^ has similarly strong association with proteins from cellular lysates, we overexpressed ^Flag^DNAJB8^WT^ or mock (eGFP) in HEK293T cells, lysed, and immunoprecipitated using the M2 anti-Flag antibody crosslinked to magnetic beads (**Figure 2A** and **Figure S1**). To minimize non-specific interactions, beads were washed well with RIPA buffer. This high detergent solution (150 mM NaCl, 50 mM Tris pH 7.5, 1% Triton X100, 0.5% sodium deoxycholate, 0.1% SDS) was originally developed to break up weak or non-specific protein-protein interactions.

**Figure 2:**
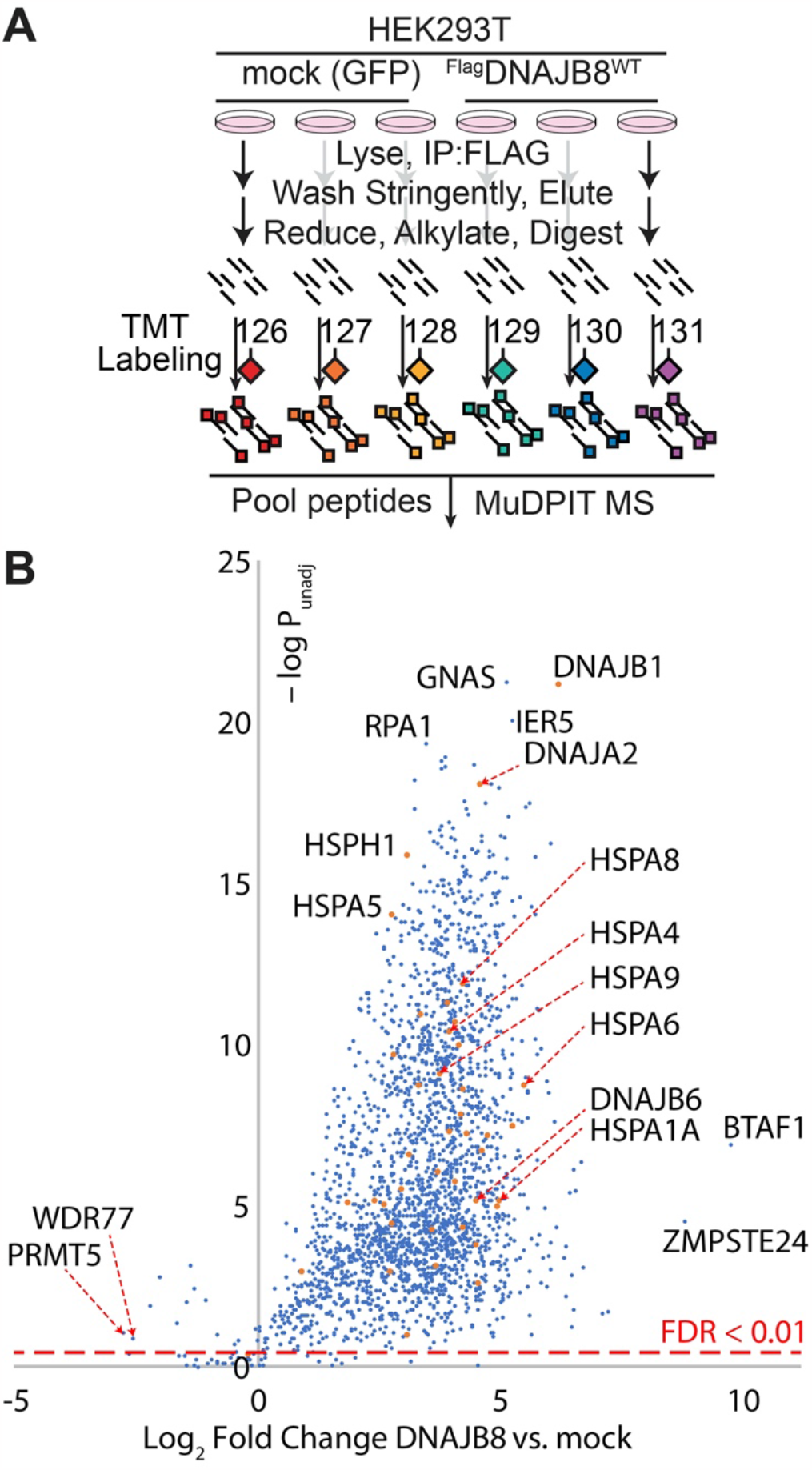
**A**. Experimental design to identify strong ^Flag^DNAJB8^WT^ interactors from HEK293T cellular lysates. Nine biological replicates, comprising three LC-MS runs, were performed. **B**. Volcano plot of proteins recovered from ^Flag^DNAJB8^WT^ immunoprecipitates. The global false discovery rate for all proteins, using Storey’s modification of Benjamini-Hochberg analysis, is 2.6%. The dashed red line on the chart shows the 1% FDR cut-off. 60 proteins that were not quantified in any mock replicate are not shown.

Tryptic digests of the eluate were isobarically labeled with TMT tags, quantified by MuDPIT LC-MS, and relative protein abundances inferred from TMT reporter ion ratios for identified peptides. Interaction significance was determined using our previously reported bait correlation method[9].

We identified 2743 proteins across the three runs, with 2623 showing preferential recovery (q-value < 0.01) in the presence of DNAJB8 (**Figure 2B** and **Table S1**). These proteins are highly enriched in RNA binding proteins (705 identified out of 1689 annotated, GO:0003723). Proteins that are selective for the *absence* of DNAJB8 include KIF11, WDR77, and PRMT5, which are well-characterized binders to anti-Flag antibodies and are prominent in control Flag immunoprecipitations in the CRAPome database[17,18]. Presumably, these proteins bind to anti-Flag antibodies that are not bound to Flag-containing protein. Piette *et al* recently reported a careful identification of human Hsp40 interactors in the cell based on AP-MS experiments and global comparison to controls[19]. They found 34 high-confidence DNAJB8 interactors, of which we identify 30 in our assay. It is important to note that their study was designed to discover specific *native interactors* of all human Hsp40s; by contrast, our assay does not require that DNAJB8 affinity only be determined for bona fide clients of the co-chaperone and we make no efforts to discriminate between native clients and the rest of the proteome. DNAJB8 co-immunoprecipitates several cytosolic Hsp70-associated proteins, including constitutive cytosolic Hsp70 HSPA8, the inducible cytosolic Hsp70s HSPA1A and HSPA6, and cytosolic Hsp70 co-chaperones STUB1, HSPA4, DNAJB1, DNAJB6, and DNAJA2. These strong associations are consistent with the canonical role of Hsp40 proteins as co-chaperones of Hsp70. In addition, the ER Hsp70 HSPA5 and mitochondrial Hsp70 HSPA9 are also recovered (alongside 183 and 112 other mitochondrial and ER proteins respectively). While mitochondrial and ER pre-proteins can interact with cytosolic chaperones prior to their trafficking or during degradation, it is likely that these associations are taking place post-lytically, and do not represent *native* clients in the cell. In the context of an assay for misfolded protein, however, post-lytic interactions serve to expand the profiling space. The large number of proteins that co-purify with DNAJB8^WT^ indicate that J-domain ablation is not necessary for the strong protein binding properties of DNAJB8.

### Influence of crosslinking and J-domain inactivation on DNAJB8 client binding

Our previous interacting protein analysis for DNAJB8^H31Q^, in the presence of the cell-penetrable crosslinker DSP, found 463 interacting proteins (using the criteria that p < 0.05, fold change > 1.2 vs. mock; 476 with q < 0.01) [9]. Of these, 251 are shared with DNAJB8^WT^ using the same criteria, while 2183 protein groups were high-confidence interactors with DNAJB8^WT^ without crosslinking but not DNAJB8^H31Q^ with crosslinking.

To better understand the role of crosslinking and J-domain inactivation in DNAJB8 interactor recovery, we performed a series of TMT-AP-MS experiments directly comparing four conditions: WT vs. H31Q, and ± crosslinker (**Figure 3A** and **Figure S2A**). For these experiments, we used the reversible crosslinker DSP. DSP is cell-penetrable and allows us to immortalize cellular interactions prior to lysis [20]. After immunoprecipitation and elution, we reverse the crosslinks with TCEP to allow peptide identification during mass spectrometry. Because crosslink yield tends to be low on a per peptide basis, we do not include DSP modification as a variable modification for peptide-spectral matching. An initial optimization found that protein recovery closely tracked DNAJB8 levels regardless of crosslinker concentration, (**Figure S2B**,**C**), so we went forward with 1 mM as the same concentration that we previously used [9]. We confirmed that DNAJB8^WT^ and DNAJB8^H31Q^ have similar expression and immunoprecipitation efficiencies (**Figure 3B** and **Figure S2D**,**E**). We expected that J-domain mutation would decrease interactions with Hsp70s, and indirectly decrease interactions with Hsp70 co-chaperones, while crosslinking would increase the recovery of most protein interactors by preventing their dissociation during lysis and washing of the beads with RIPA.

**Figure 3:**
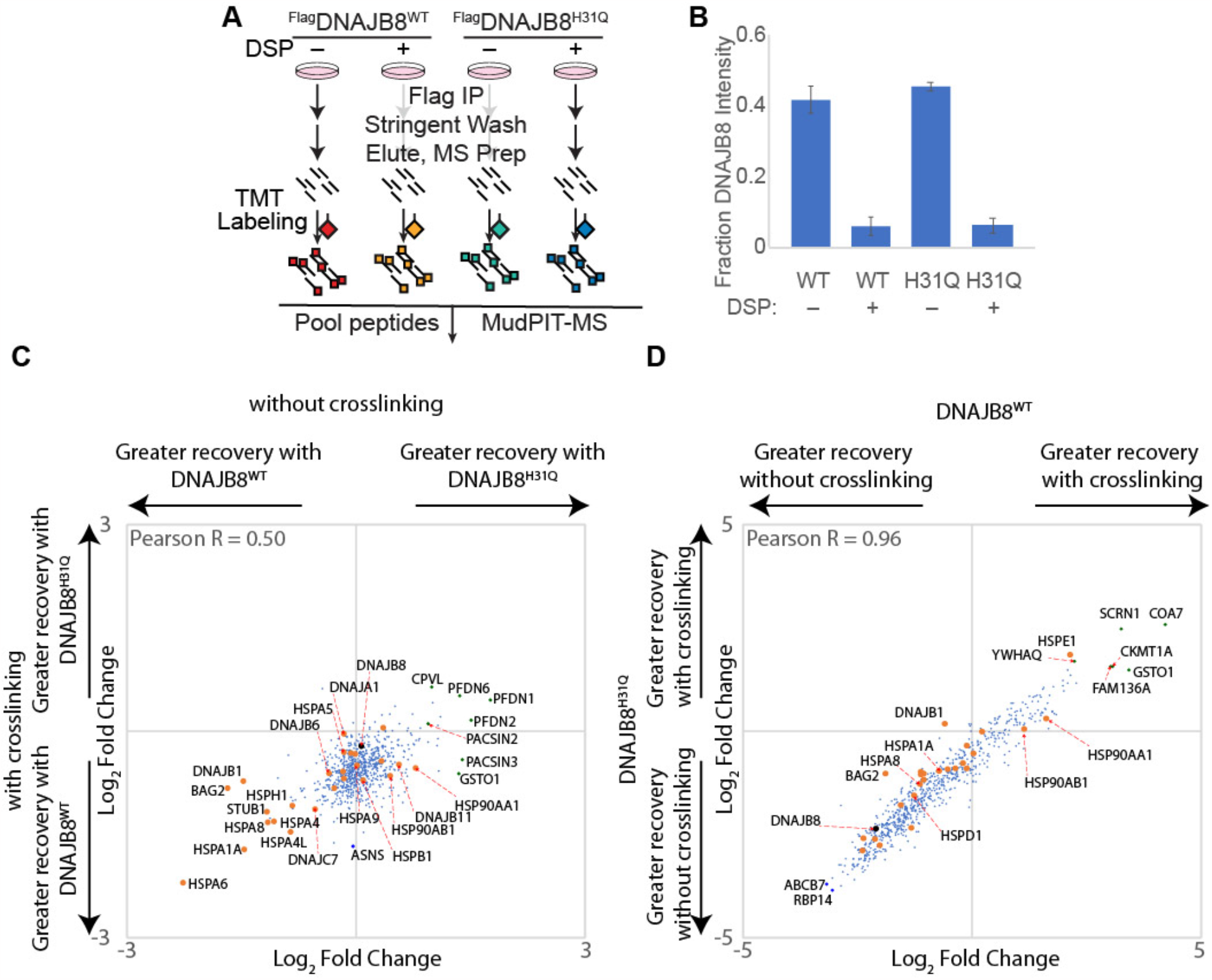
Effect of J-domain integrity and cellular crosslinking on interactor recovery with DNAJB8. **A**. Schematic of immunoprecipitation. **B**. Recovery of DNAJB8 from each condition. Error bars represent standard deviation (n = 3). **C, D**. Relative association of interactors. Hsp70 chaperones and Hsp70-associated co-chaperones are shown in orange. Three biological replicates, comprising three LC-MS runs, were performed.

J-domain mutation has only a modest effect on the interactor profile of DNAJB8 (**Figure 3C, Figure S3**, and **Table S2**). As expected, cytosolic Hsp70 chaperones and associated co-chaperones have higher affinity for DNAJB8^WT^ than for DNAJB8^H31Q^, reflecting the role of the J-domain in mediating their binding. This is true both in the presence and absence of crosslinking. By contrast, the ER and mitochondrial Hsp70s (HSPA5, HSPA9), have similar affinity for both DNAJB8s (**Figure S3B**), suggesting that the J-domain is not mediating interactions with Hsp70s that DNAJB8 would not encounter during normal expression in the cell. The proteins with the strongest preference for DNAJB8^H31Q^ compared to DNAJB8^WT^ are the prefoldin ? subunits (PFDN1, PFDN2, PFDN6) (**Figure S3B**). No such preference is found for components of the prominent prefoldin-associated complexes TRiC and RNA polymerase II, suggesting that the recognized prefoldin subunits are not actively engaged in those canonical prefoldin complexes [21]. Prefoldin subunits have also been found to act as holdase chaperones separately from their interactions with the TriC complex[22]. It could be that prefoldin is recovered through a tertiary interaction, mediated by misfolded proteins that are bound jointly to DNAJB8 and prefoldin. In this case, the relatively lower recovery with DNAJB^WT^ as opposed to DNAJB8^H31Q^ could be ascribed to competition with Hsp70 that is recruited by the DNAJB8 J-domain. Alternatively, there could be a misfolded population of DNAJB8^H31Q^ that is preferentially and stably bound by prefoldin subunits, but not other cellular chaperones.

As expected, crosslinking sharply decreases DNAJB8 levels in the lysate as well as the recovered fraction, by > 80% (**Figure 3B**). While this decrease could be due to DSP modification of the lysine-rich Flag tag, the total protein concentration as measured by Bradford assay decreased by 48% ± 2% with DSP crosslinking, indicating that decreased protein solubility meaningfully contributes. Interestingly, crosslinking increases immunoprecipitation efficiency, perhaps by rendering any large DNAJB8-containing complexes insoluble (**Figure S2D**,**E**). The effect of crosslinking on interactor recovery is almost identical between the DNAJB8 baits, with a Pearson correlation of 0.96 between the two profiles (**Figure 3D, Figure S3**, and **Table S1**). Crosslinking sharply decreases recovery of DNAJB8, and although it increases the recovery of interactors *relative to* DNAJB8, it is not enough to offset the decrease in DNAJB8 bait for most interactors. Exceptions include HSPE1 and a few 14-3-3 proteins, particularly 14-3-38/YWHAQ (**Figure S3B**). In summary, the primary consequence of DNAJB8 J-domain inactivation is to decrease association to Hsp70 family chaperones and co-chaperones, while crosslinking decreases DNAJB8 recovery so drastically as to eliminate any benefit from greater protein recovery.

### Interactor Recovery by DNAJB1 Immunoprecipitation

Class B Hsp40s are distinguished by an N-terminal J-domain, a glycine/phenylalanine rich domain, two beta-barrel domains, and a C-terminal dimerization domain [6]. For cytosolic Class B Hsp40s, the first beta-barrel includes a weak Hsp70 binding site which is important for client transfer[23]. This class can be further divided into the two phylogenetic trees [24]. In one branch, DNAJB6 and DNAJB8 feature a unique serine/threonine rich region. This region is implicated in their remarkable ATP-independent holdase activity that substoichiometrically inhibits aggregation of some proteins[25–28], while still requiring Hsp70 to inhibit aggregation of other susbtrates[29]. The most abundant Hsp40 of the other branch is DNAJB1 which is homologous to yeast Sis1. DNAJB1 primarily forms dimers and monomers, and has been found to demonstrate different substrate specificity in cellular and in vitro assays as opposed to DNAJB6 and DNAJB8 [25,28,30].

We characterized the interaction networks of ^Flag^DNAJB1^WT^ (in the absence of crosslinking), and ^Flag^DNAJB1^H32Q^ (in the presence of crosslinking), to determine profiles of potential interactors (**Figure 4A**,**B, Figure S4A**, and **Table S3**). While these conditions do not allow direct comparison to each other, they do allow comparisons to the equivalent DNAJB8 experiments of **Figure 2** and reference 8. The crosslinking concentration was based on optimization by TMT-AP-MS at varying concentrations of DNAJB1^H32Q^ (**Figure S4B**,**C**). Although median reporter ion intensities only change modestly with varying [DSP], there was a slight maximum at 0.25 mM DSP. We also found that extending the immunoprecipitation to 3 days increased DNAJB1 recovery (**Figure S4D**,**E**). DNAJB1^WT^ without crosslinking co-immunoprecipitates fewer proteins than DNAJB8^WT^ (**Figure 2**), and the strongest interactors are almost entirely Hsp70s or their associated co-chaperones (**Figure 4A**). 14/14 of the native interactors from Piette *et al*[19] were found, though two (PYCR3, RPS10) did not show meaningful selectivity in our experiment between the presence or absence of DNAJB1^WT^. 15/33 of Bioplex interactors[31,32] (from HEK293) were identified, of which 4 (PYCR3, HDLBP, MAP2K2, and DIS3) did not show meaningful selectivity. These proteins participate in extensive interaction networks per Bioplex data, and they might lose affinity to DNAJB1 when these networks are disrupted by highly stringent RIPA buffer. As with DNAJB8^WT^, common anti-Flag-binding contaminants are depleted in the ^Flag^DNAJB1^WT^ immunoprecipitates. While DNAJB1^H32Q^ with crosslinking robustly recovers a larger proteome, the strongest interactions are still dominated by Hsp70 and Hsp70-associated chaperones. This is at first surprising, given the J-domain ablation. While the Hsp70 family proteins could be associating with DNAJB1^H32Q^ as clients, or as chaperones for misfolded DNAJB1^H32Q^, the most likely explanation is that DNAJB1^H32Q^ is forming heterodimers with endogenous DNAJB1^WT^ or other endogenous J-domain proteins. Such heterodimers have been observed for most Class A and Class B DNAJ proteins[33,34]. Still, DNAJB1^H32Q^ does interact with some proteins that were not recovered by either DNAJB8^WT^ or DNAJB8^H31Q^, suggesting that it could potentially be useful for extending the space of proteins that are sampled by Hsp40 affinity profiling (**Figure S5**).

**Figure 4.**
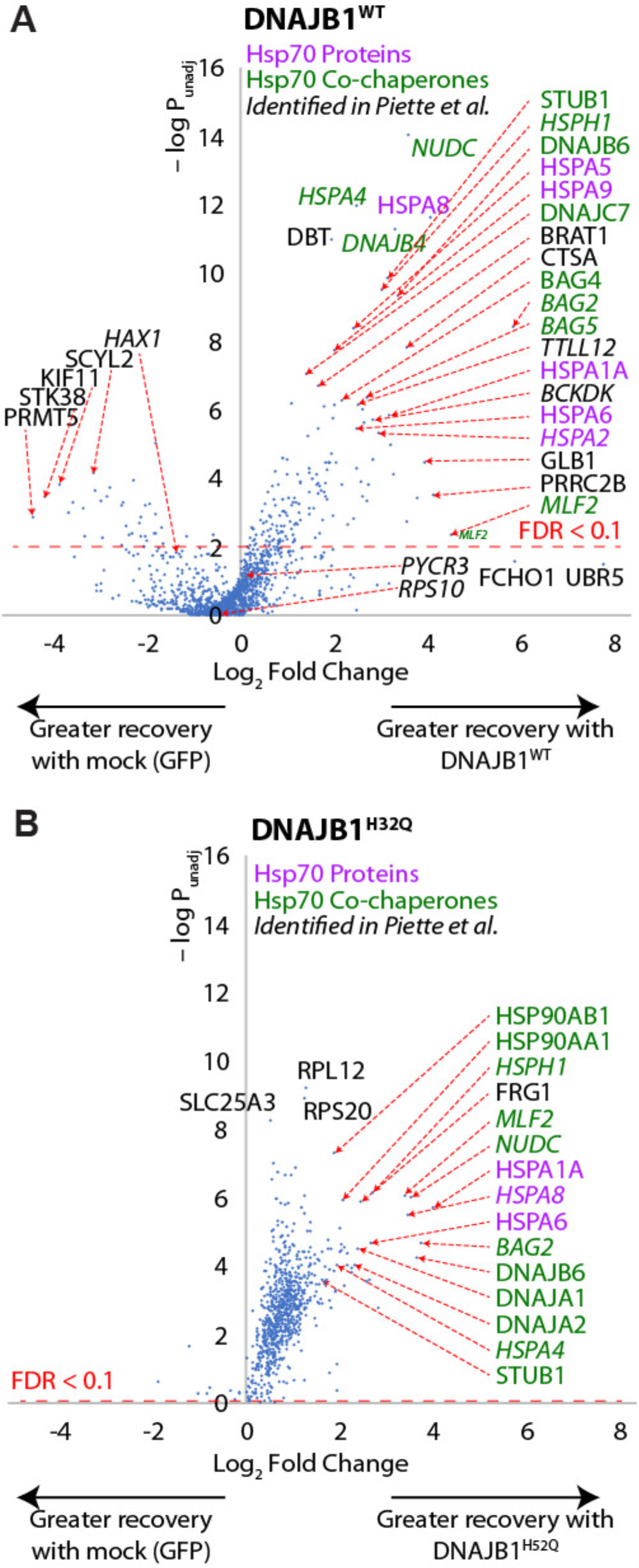
**A**. Proteins recovered by immunoprecipitation of ^Flag^DNAJB1^WT^ without crosslinking (comparable to **Figure 2B**) and **B**. Proteins recovered by immunoprecipitation of ^Flag^DNAJB1^H32Q^ with crosslinking (comparable to ref 8). Mock cells are transfected with eGFP. The experimental strategy is similar to that from **Figure 2A**, except that the DNAJB1^H32Q^ AP-MS was preceded by cellular crosslinking with 250 µM DSP. Nine biological replicates, comprising three LC-MS runs, were performed for DNAJB1^WT^, while six biological replicates, comprising two LC-MS runs, were performed DNAJB1^H32Q^. Hsp70 proteins are indicated in purple, Hsp70-associated co-chaperones are in green, and protein identified from Piette *et al*[19] are italicized. The p-value cut-off corresponding to a 10% False Discovery Rate is indicated. The global false discovery rate for all proteins from Storey’s modification of Benjamini-Hochberg analysis is 73% for DNAJB1^WT^ and 5.9% for DNAJB1^H32Q^.

To better understand the role that the J-domain has in DNAJB1 interactor recovery, we directly compared interactor recovery for DNAJB1^WT^ vs. DNAJB1^H32Q^ in both the absence and presence of crosslinking; this is similar to the experiment described in **Figure 3A** for DNAJB8. In contrast to DNAJB8, there is far lower expression and hence recovery of DNAJB1^H32Q^ as opposed to DNAJB1^WT^ (**Figure 5A, Figure S6A**, and **Table S4**). Similar loss of DNAJB1 for both WT and H32Q is observed with crosslinking. For both crosslinking and J-domain inactivation, however, the loss of bait is offset by a corresponding increase in protein recovery relative to DNAJB1 levels, such that overall interactor recoveries are similar across all four conditions (**Figure 5B**,**C** and **Figure S6B**). As seen in **Figure 4B**, DNAJB1^H32Q^ particularly enriches the inducible cytosolic Hsp70s HSPA6 and HSPA1A, despite lacking an active J-domain (**Figure 5B** and **Figure S6B**). The WT protein, on the other hand, preferentially interacts with BCKDK and TTLL12 (**Figure 5B and Figure S6B**). Crosslinking is necessary for recovery of 14-3-3 proteins and Hsp90s, but lowers recovery of BCKDK (**Figure 5C and Figure S6B**). The low overall protein recovery made us consider that perhaps RIPA washing was responsible for removing interactors. Hence, we performed an identical set of experiments using a gentle lysis and wash buffer (0.1% Triton X100 in TBS). Nearly identical results were obtained, except that overall protein identifications dropped to only 284 proteins (**Figure S7**). This decrease in protein recovery could be due to the low detergent buffer leading to less efficient lysis. While recovery in the DNAJB1^WT^ is unaffected, gentle washing increases protein recovery with DNAJB1^H32Q^ in the absence of crosslinking as opposed to in the presence of crosslinking. Overall, J-domain inactivation and crosslinking are, both individually and combined, effective approaches to increase interactor stoichiometry on DNAJB1. However, the low recovery of DNAJB1 itself under both of these conditions offsets the greater interactor stoichiometry. This challenge would have to be overcome to make DNAJB1 a useful tool for separation of misfolded cellular proteins.

**Figure 5:**
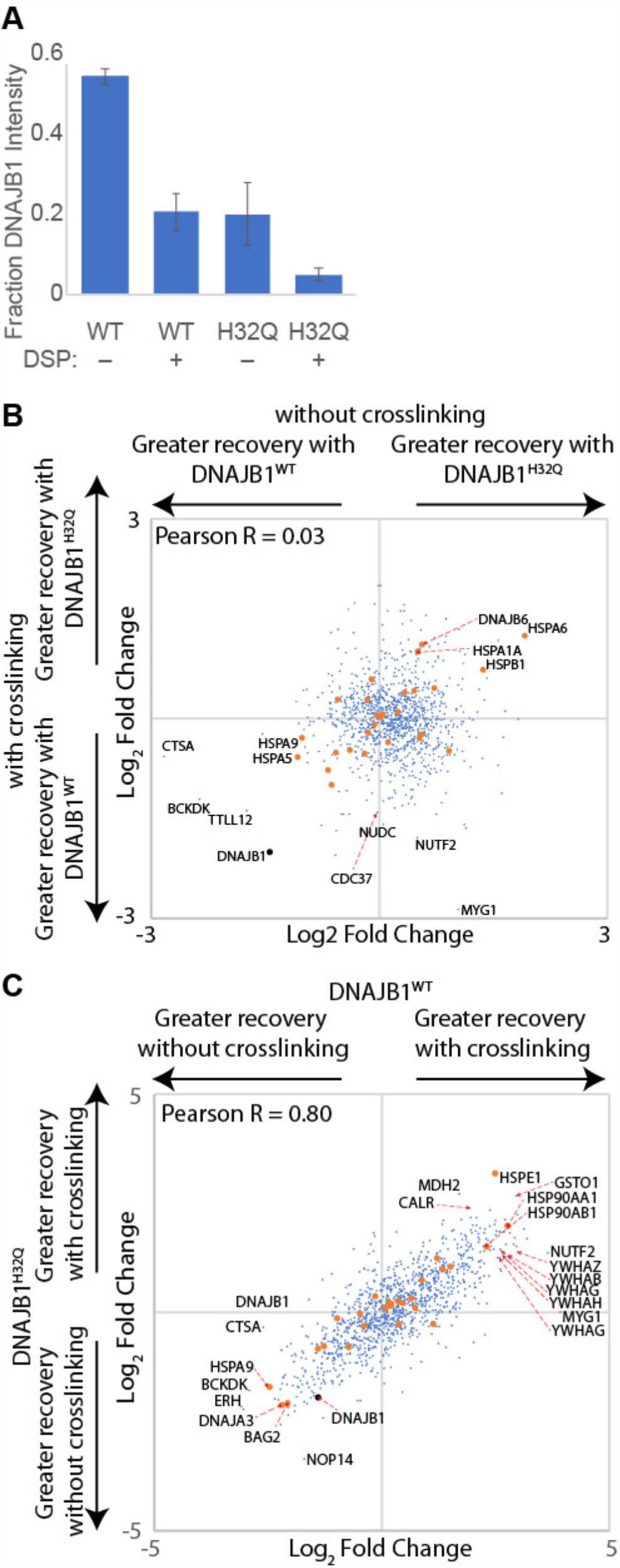
Effect of J-domain integrity and cellular crosslinking on interactor recovery with DNAJB1. **A**. Schematic of immunoprecipitation. **B**. Recovery of DNAJB1 from each condition. Error bars represent standard deviation (n = 3). **C, D**.. Relative association of interactors. Hsp70 chaperones and Hsp70-associated co-chaperones are shown in orange. Three biological replicates, comprising three LC-MS runs, were performed.

### DNAJB8 demonstrates excellent client recovery, both with and without J-domain inactivation

The binding is strong even without crosslinking and following stringent washing, allowing elimination of most non-specific interactors. We considered how DNAJB8^WT^ and DNAJB8^H31Q^ compare with regards to their ability to identify changes in protein stability following a stress. We subjected ^Flag^DNAJB8^H31Q^-expressing HEK293T cells to mild heat stress for 30 min., followed by immediate lysis and anti-Flag immunoprecipitation (**Figure 6A**). A condition with mock transfection (eGFP) under 47 °C heat shock was included to control for misfolding-induced affinity for beads. The short treatment was chosen to minimize transcriptional/translational remodeling of the cell due to induction of the Heat Shock Response[35,36]. The relative recovery of proteins was determined by MuDPIT LC-MS with isobaric TMT labeling, and integrated reporter ion ratios normalized to the amount of DNAJB8 bait. Proteins that showed less than 2-fold selectivity for the presence of DNAJB8 were excluded from further analysis (34 protein groups, dominated by the usual anti-Flag binding proteins discussed above), leaving 989 protein groups identified and quantified from all three runs. Consistent with the validated ability of DNAJB8^H31Q^ affinity to serve as a proxy for protein stability, we find that for about 80% of the proteome the Hsp40 affinity monotonically increases as the temperature increases from 37 °C to 45 °C, with plateauing or a slight drop-off at 47 °C (**Figure 6B**,**C, Figure S8A**,**B**, and **Table S5**). For proteins that do not exhibit this trend, there is a tendency for Hsp40 affinity to increase from 37 °C to 43 °C, followed by a decrease in protein recovery. Although we did not collect sufficient data to estimate transition temperatures, we can estimate the sensitivity of Hsp40 affinity to temperature by taking the response factor (slope). We compared these slopes to published aggregation temperatures in HEK293T cells as determined by CETSA[37] and melting temperatures in HeLa cells determined from limited proteolysis[38] (**Figure S8E**). No correlation is found (643 protein groups found in both data sets, R = 0.0008), as has been seen in other studies comparing relative destabilization using different methods [39]. We then performed a similar experiment with ^Flag^DNAJB8^WT^, now replacing the mock condition with another temperature as little protein binding recovery had been seen in the mock condition (**Figure 6D**). Surprisingly, we see only modest changes in DNAJB8^WT^ affinity with increasing temperature (**Figure 6E**,**F, Figure S8C**,**D**, and **Table S5**). Although a few proteins show increased association, most show no change. Hence, while DNAJB8^H31Q^ is effective for recovering proteins that are destabilized by stress, DNAJB8^WT^ does not show the same capability.

**Figure 6.**
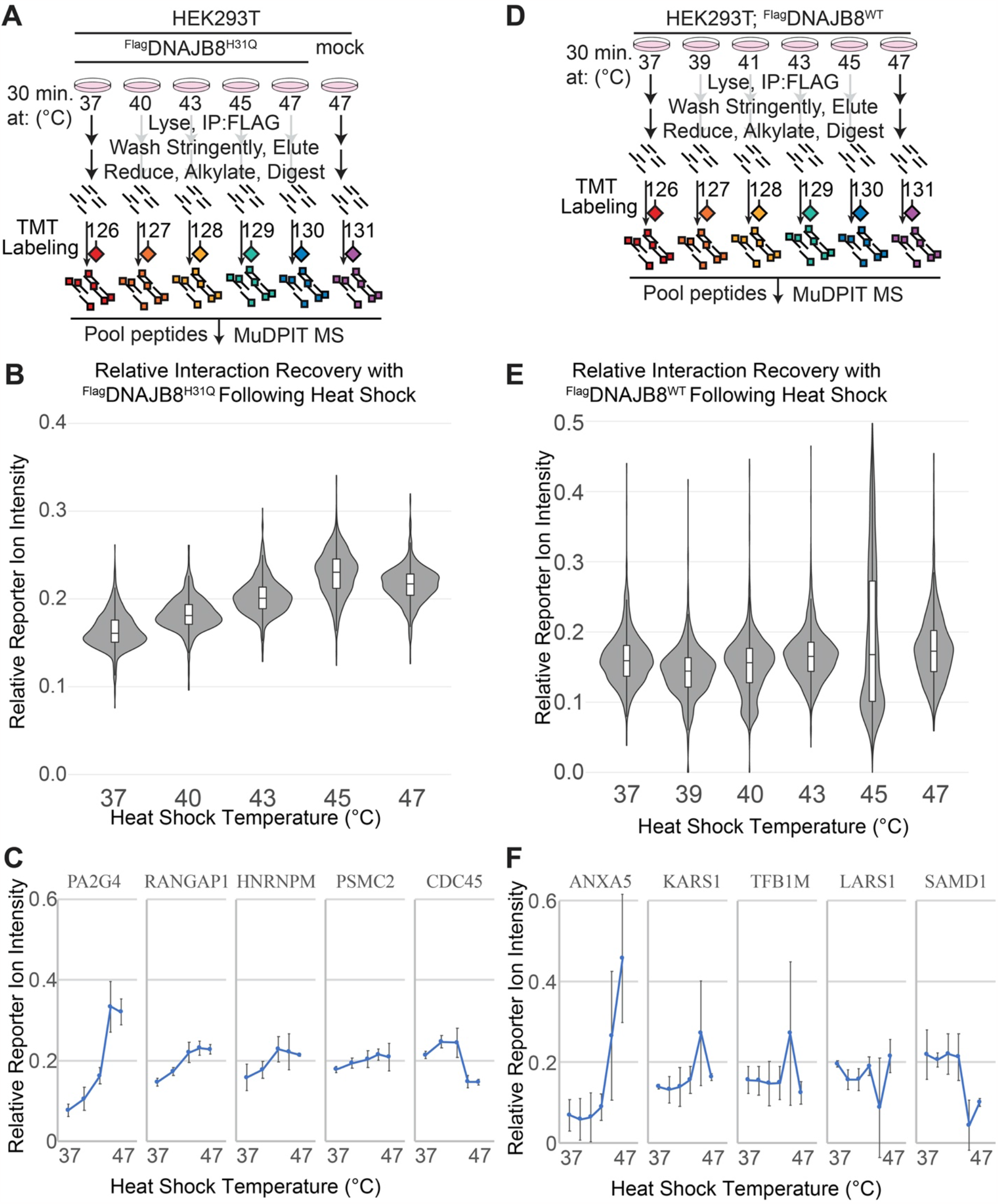
**A**. Schematic describing the experiment profiling DNAJB8^H31Q^ affinity following heat shock. **B**. Violin and bar plots for all quantified interacting proteins (n = 989). **C**. Changes in reporter ion intensity for proteins with the highest, 25%, median, 75%, and lowest response with respect to temperature. Error bars represent standard deviation (n = 3). **D**. Schematic describing the experiment profiling DNAJB8^WT^ affinity following heat shock. **E**. Violin and bar plots for all quantified interacting proteins (n = 1541). **F**. Changes in reporter ion intensity for proteins with the highest, 25%, median, 75%, and lowest response with respect to temperature. Error bars represent standard deviation (n = 3).

## DISCUSSION

We previously demonstrated that DNAJB8^H31Q^ is effective for profiling the misfoldome in response to cellular stress[3], due to seemingly irreversible binding to misfolded proteins. DNAJB1 is one of the best studied Hsp40s due to its high concentration, promiscuous activity, selectivity for misfolded protein, and high similarity to yeast Sis1[6,23,40,41], leading us to consider whether DNAJB1 could be similarly used to recover misfolded proteins. We find that DNAJB1^WT^ preferentially associates with Hsp70 family members, pulling down a far less diverse proteome than DNAJB8 (**Figure S5**), consistent with its primarily ATP (and presumably Hsp70)-dependent biological function. While J-domain inactivation and crosslinking both help increase the relative protein load on DNAJB1, they decrease bait recovery so much that any benefit is offset. Crosslinkers vary in the functional groups that they target, in their solubility, and in the length between the reactive functional groups. Expanding the range of crosslinkers used beyond DSP would be useful in case other crosslinkers provide improved performance. Furthermore, other Class B Hsp40 chaperones with substantial ATP-independent activity such as DNAJB6 or DNAJB2a might also serve as factors to expand the client profile accessible through DNAJB8^H31Q^. It is important to stress that there is no reason to believe that the proteins recognized through this AP-MS assay reflect *native* cellular clients of Hsp40’s. Rather, many of the interactors, such as those to mitochondrial and secretory proteins, are taking place post-lytically.

Given that both DNAJB8^WT^ and DNAJB8^H31Q^ strongly bind a large proteome, we expected that heat shock would have a similar effect on protein affinity for both chaperones. This was not the case, as heat shock increases client protein binding to DNAJB8^H31Q^ and not DNAJB8^WT^ (**Figure 6**). The small change in client affinity for DNAJB8^WT^, as opposed to DNAJB8^H31Q^, could be due to the enhanced proteostasis activity following stress. If increased protein association with DNAJB8^WT^ during heat shock also increases the flux of hand-off to Hsp70, then the steady state association of client proteins will only modestly change. DNAJB8^H31Q^, with its impaired hand-off to Hsp70, will have increased binding with the newly misfolded proteins during heat shock, as we observe. One interesting unaddressed question is the role of DNJAB8 oligomerization. DNAJB8 and the structurally similar DNAJB6 form a distribution of oligomeric, dimeric, and monomeric stables, while DNAJB1 forms dimers [26,42,43].

In summary, DNAJB1 and DNAJB8 have been evaluated for their ability to profile proteins as part of an Hsp40 affinity assay. We find that DNAJB8^H31Q^ without crosslinking is the most effective approach and demonstrate that mild heat shock leads to generally monotonic increased affinity of clients with DNAJB8^H31Q^.

## Supporting Information Available

Supporting Figures S1 through S8, providing silver stains, Western blots, and further analysis of proteomic data (PDF)

- Table S1 providing integrated TMT reporter ions and analysis for DNAJB8^WT^ AP-MS interactors.
- Table S2 providing integrated TMT reporter ions and analysis for AP-MS comparing DNAJB8^WT^ and DNAJB8^H31Q^ with or without crosslinking.
- Table S3 providing integrated TMT reporter ions and analysis for DNAJB1^WT^ and DNAJB1^H32Q^ AP-MS interactors.
- Table S4 providing integrated TMT reporter ions and analysis for AP-MS comparing DNAJB1^WT^ and DNAJB1^H32Q^ with or without crosslinking.
- Table S5 providing integrated TMT reporter ions and analysis for the effect of heat shock treatment on DNAJB1^WT^ and DNAJB1^H32Q^ AP-MS.

The mass spectrometry proteomics data and associated results files have been deposited to the ProteomeXchange Consortium via the PRIDE partner repository with the dataset identifier PXD030633.

## ACKNOWLEDGEMENTS

eGFP.pDEST30 plasmid was generously shared by R.L. Wiseman. This work was supported by a Society of Analytical Chemists of Pittsburgh Starter Grant (J.C.G.) and the University of California, Riverside.

