## Supplemental Information for "Factors Affecting Protein Recovery During Hsp40 Affinity Profiling"

**Department of Chemistry, University of California, Riverside, CA 92521**

*Joseph C. Genereux,

| Page | Contents |
| --- | --- |
| S-2 | Table of Contents |
| S-3 | Supplemental Figures |
| S-12 | Supplemental References |


Supplemental Tables 1-5 are provided as external files.

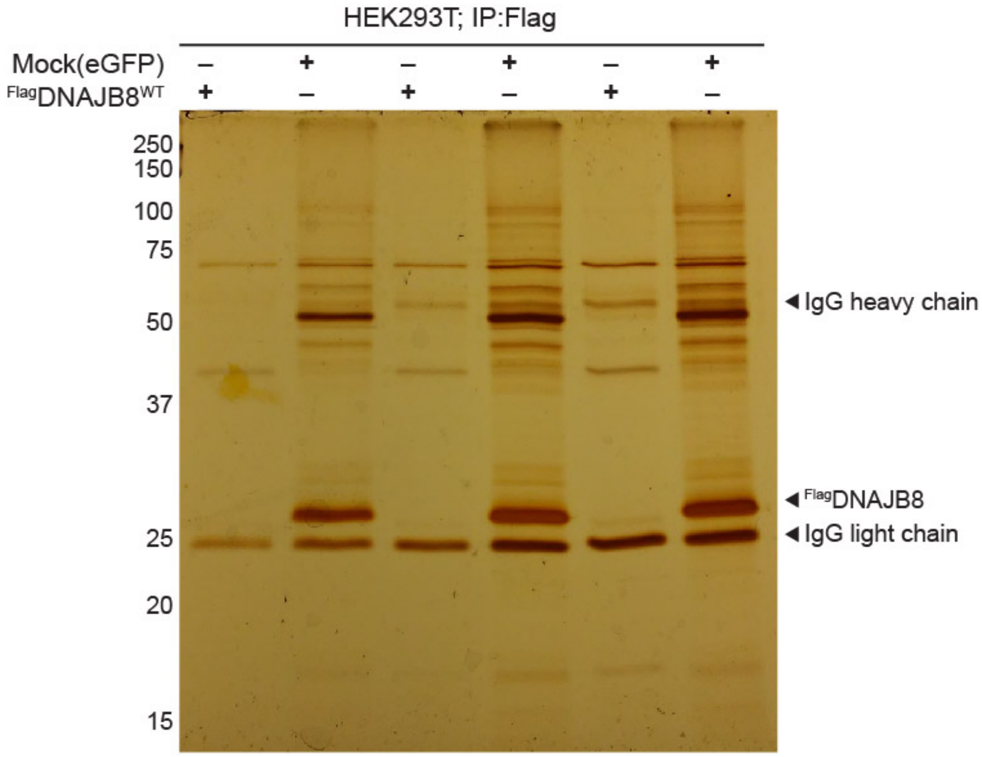


**Figure S1**. Representative Silver Stain of SDS-PAGE separated eluates associated with **Figure 2**. Based on spectral counts, the intense interactors at 50 kDa are likely tubulins, while the non-specific band at 72 kDa is likely PRMT5.

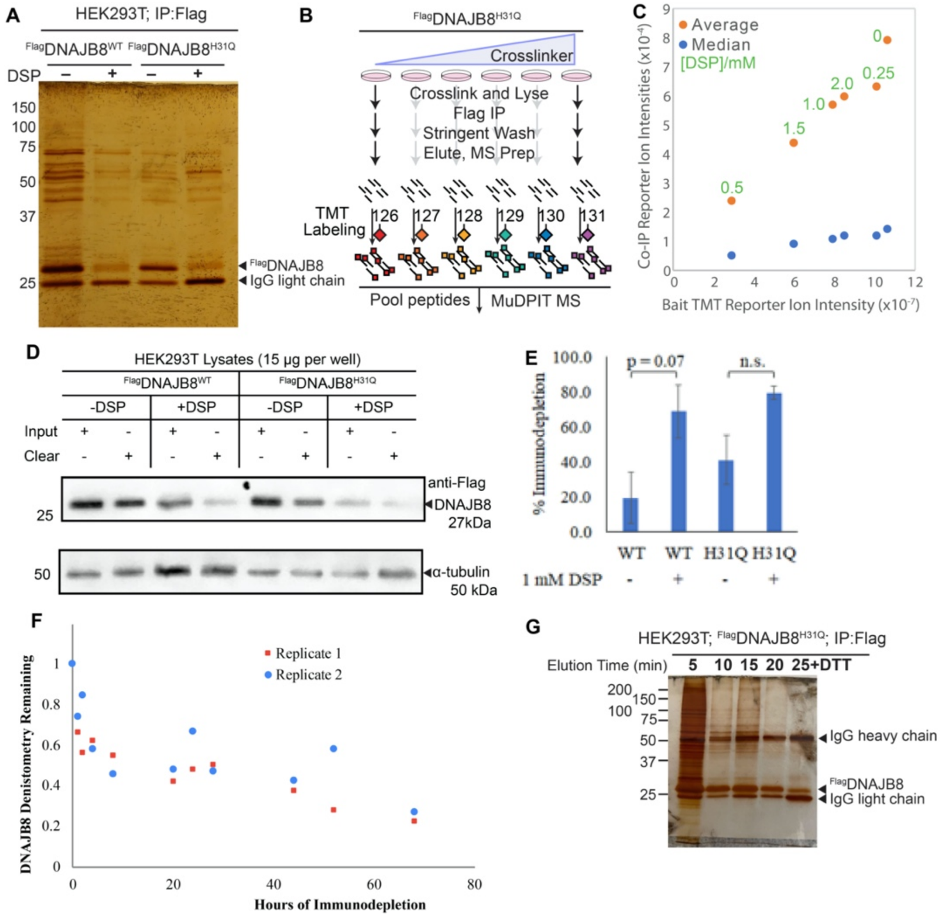


**Figure S2**. **A**. Representative Silver Stain of SDS-PAGE separated eluates associated with **Figure 3**. **B**. Schematic for crosslinking optimization experiment for DNAJB8$^{H31Q}$ recovery. C. Average and median integrated (protein group level) reporter ion intensities plotted against the integrated DNAJB8$^{H31Q}$ reporter ion intensities. The amount of crosslinker used is indicated next to the data points. Although the effect of crosslinker is not monotonic, reflecting variability in cell growth, transfection, and crosslinking efficiency between replicates, the relationship between bait intensities and interactor intensities is monotonic, reflecting that crosslinking affects interactor recovery to the extent that it decreases the amount of available protein. **D**. Representative immunoblot of SDS-PAGE separated lysates prior to and following DNAJB8 immunodepletion. **E**. Quantification of **S2D**. Error bars represent standard deviation from 3 replicates. p values were determined from a two-tailed Student's t test with a 0.1 threshold for significance. **F**. Timecourse of immunodepletion for $^{Flag}$DNAJB8$^{H31Q}$ without crosslinking. Lysate samples from HEK293T cells overexpressing $^{Flag}$DNAJB8$^{H31Q}$ were incubated on anti-Flag beads and aliquots removed at the indicated timepoint. DNAJB8 levels were determined by Western Blotting with anti-FLAG and densitometry measurement. **G**. Timecourse of elution of $^{Flag}$DNAJB8$^{H31Q}$ and associated proteins from beads. Beads were boiled in Laemmli 6X concentrate with the liquid removed and fresh Laemmli 6X concentrate added at 5 min. increments. The final elution was performed in the presence of 100 mM DTT. It is clear that nearly all protein is recovered from a 5 min. incubation.

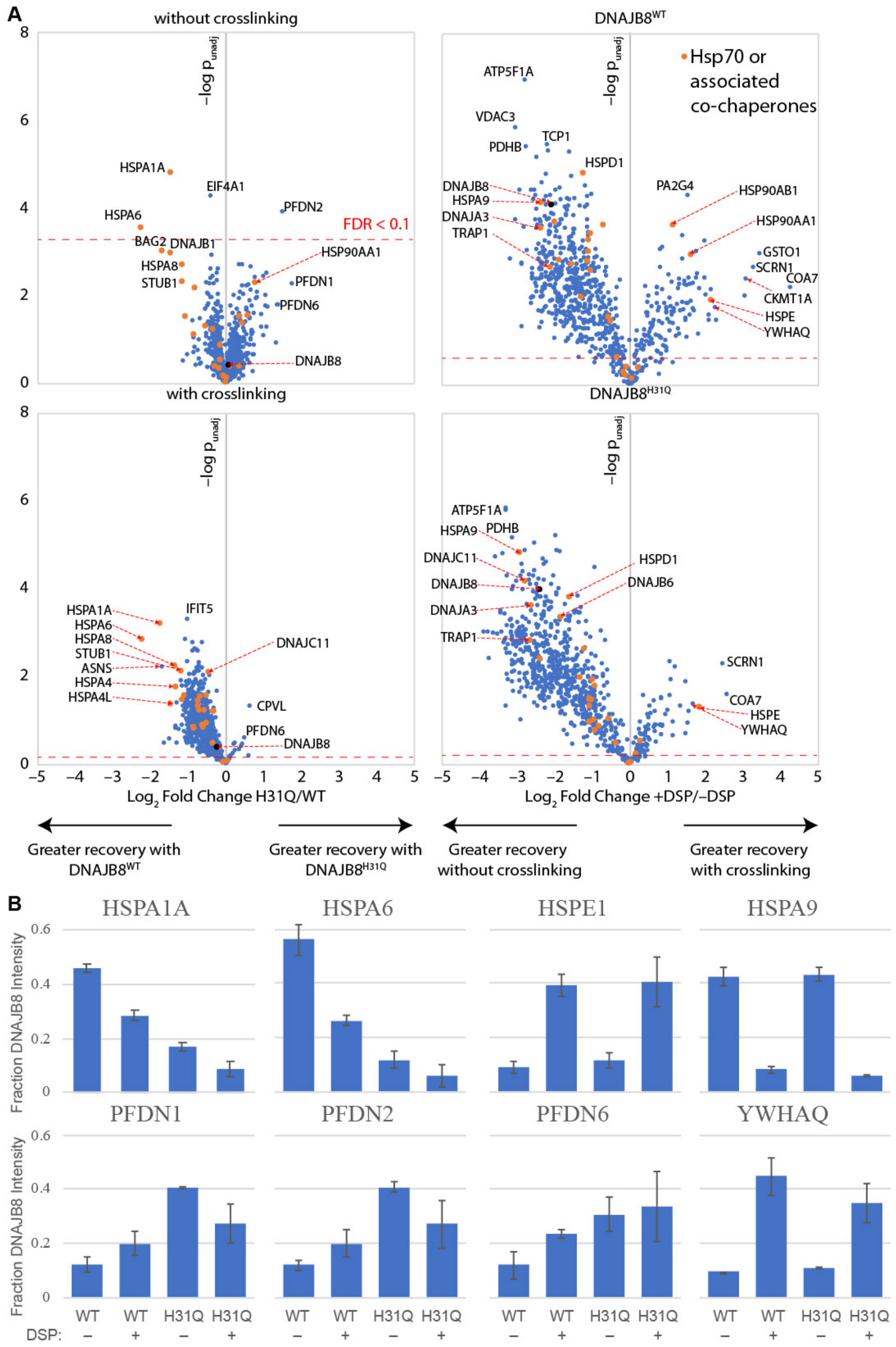


**Figure S3. A**. Volcano plots corresponding to the protein interactor changes depicted in **Figure 3C,D**. The lower threshold for a FDR < 0.1, as determined by Storey's modification of the Benjamini-Hochberg method, is indicated by the red dashed line. **B**. Relative recovery of indicated interactors across conditions described in **Figure S3A**. Error bars represent standard deviation (n = 3).

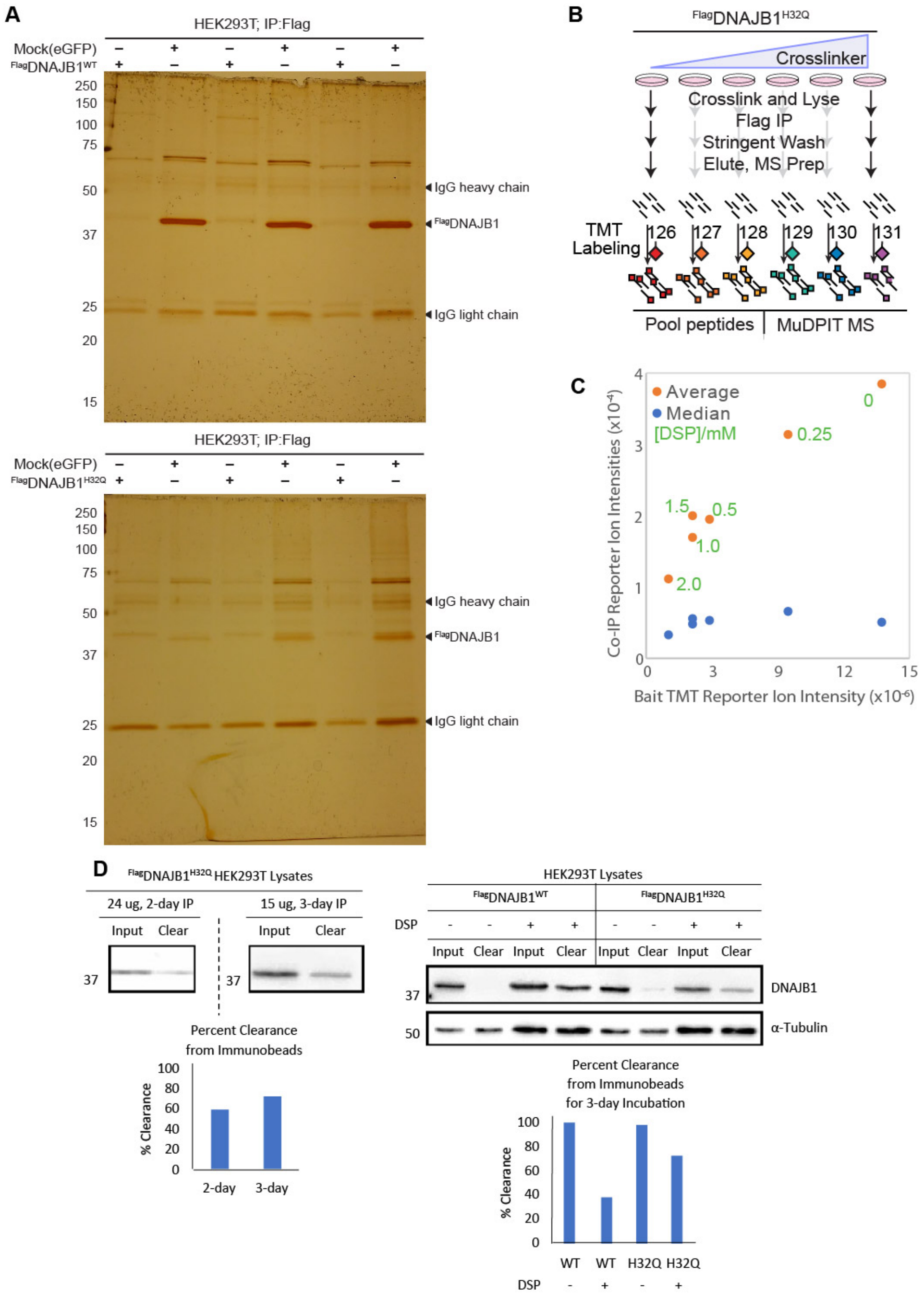


**Figure S4. A**. Representative Silver Stains of SDS-PAGE separated eluates associated with **Figure 4**. **B**. Schematic for crosslinking optimization experiment for DNAJB1$^{H32Q}$ recovery. **C**. Average and median integrated (protein group level) reporter ion intensities plotted against the integrated DNAJB1$^{H32Q}$ reporter ion intensities. The amount of crosslinker used is indicated next to the data points. Although the effect of crosslinker is not monotonic, reflecting variability in cell growth, transfection, and crosslinking efficiency between replicates, the relationship between bait intensities and interactor intensities is monotonic, reflecting that crosslinking affects interactor recovery to the extent that it decreases the amount of available protein. **D**. Representative immunoblots of SDS-PAGE separated lysates prior to and following DNAJB1 immunodepletion, with accompanying quantifications.

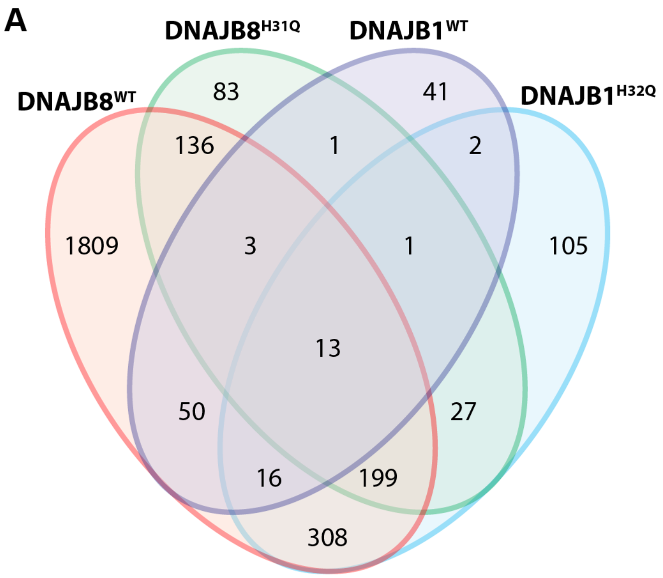


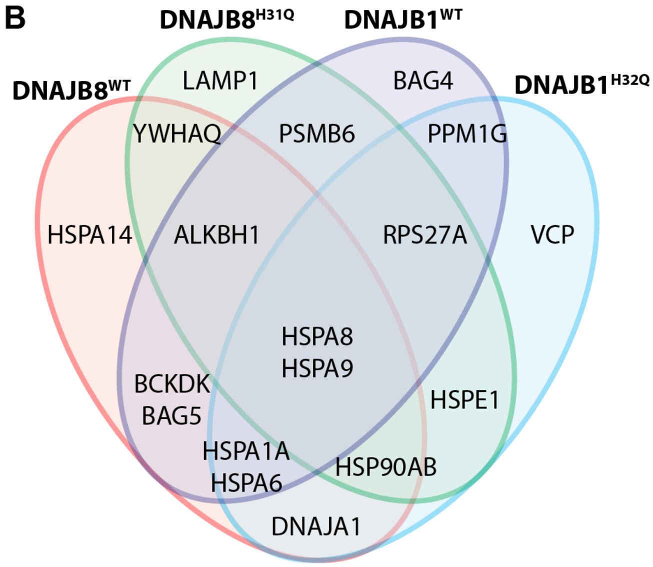


**Figure S5**. **A**. Venn diagram comparing protein interactors of DNAJB8$^{WT}$, DNAJB1$^{WT}$, DNAJB8$^{H31Q}$ (with crosslinking), and DNAJB1$^{H32Q}$ (with crosslinking), according to the threshold that fold change vs. mock is $\geq 1.2$ and that the unadjusted p-value is below 0.05. **B.** Examples of proteins in each classification.

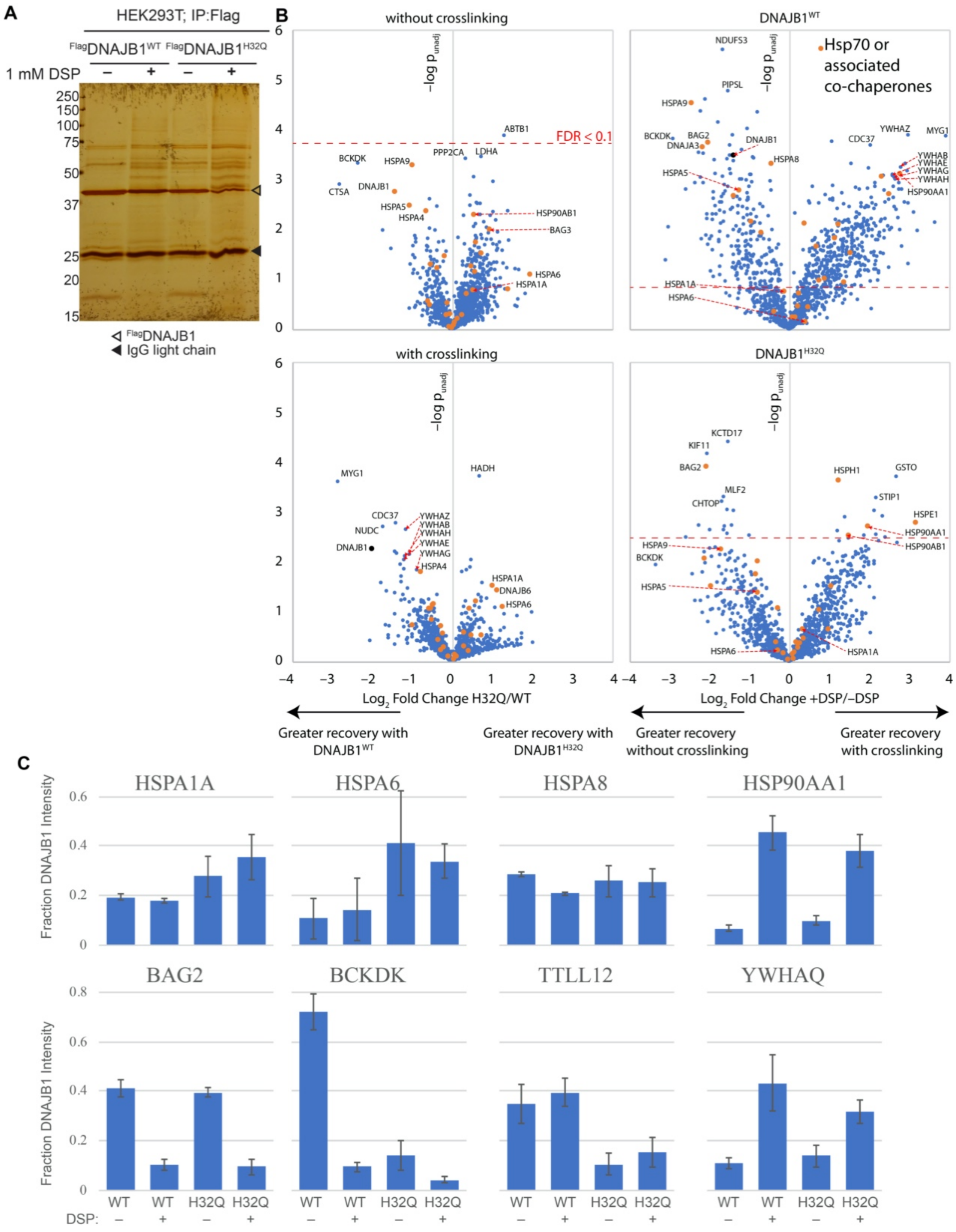


**Figure S6**. **A**. Representative Silver Stains of SDS-PAGE separated eluates associated with **Figure 5**. **B**. Volcano plots corresponding to the protein interactor changes depicted in **Figure 5B**. **C**. Relative recovery of indicated interactors across conditions as described in **Figure 5A**. Error bars represent standard deviation (n = 3).

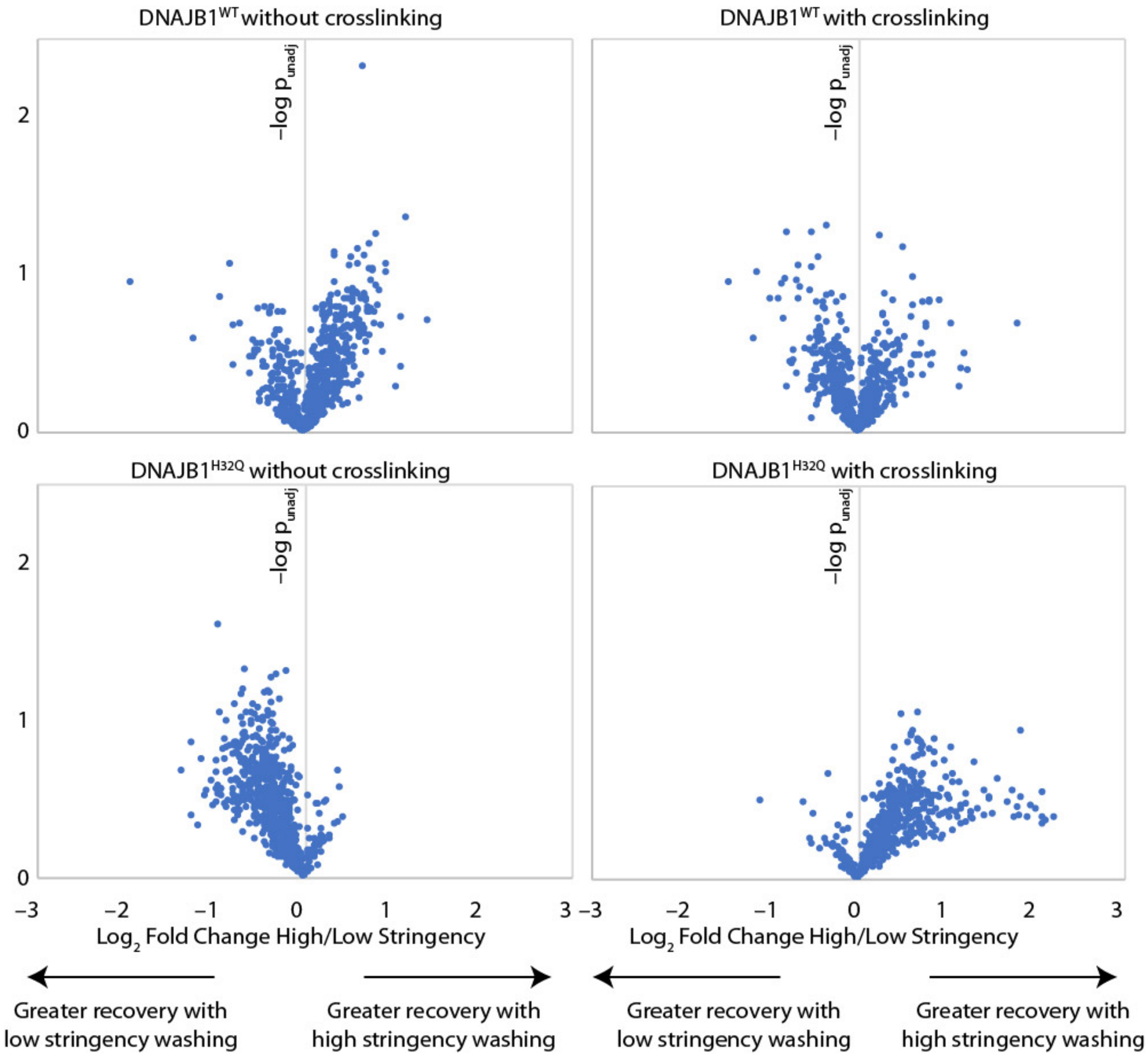


**Figure S7**. Volcano plots illustrating how low stringency (0.1% Triton X100 in TBS) vs. high stringency (RIPA buffer) lysis and washing affects protein recovery. 4-plex experiments were ran identically to as described in Figure 5A, except that instead of RIPA buffer, 0.1% Triton X100 in TBS was used for gentle lysis and bead washing to allow weaker protein-protein interactions to persist. Protein recovery is compared between high stringency and low stringency conditions for each combination of {DNAJB1$^{WT}$, DNAJB1$^{H32Q}$}⊗{–DSP crosslinking, + DSP crosslinking}. No differences between low and high stringency washing are significant beyond the 1% FDR threshold; the lowest q-value observed was above 0.7.

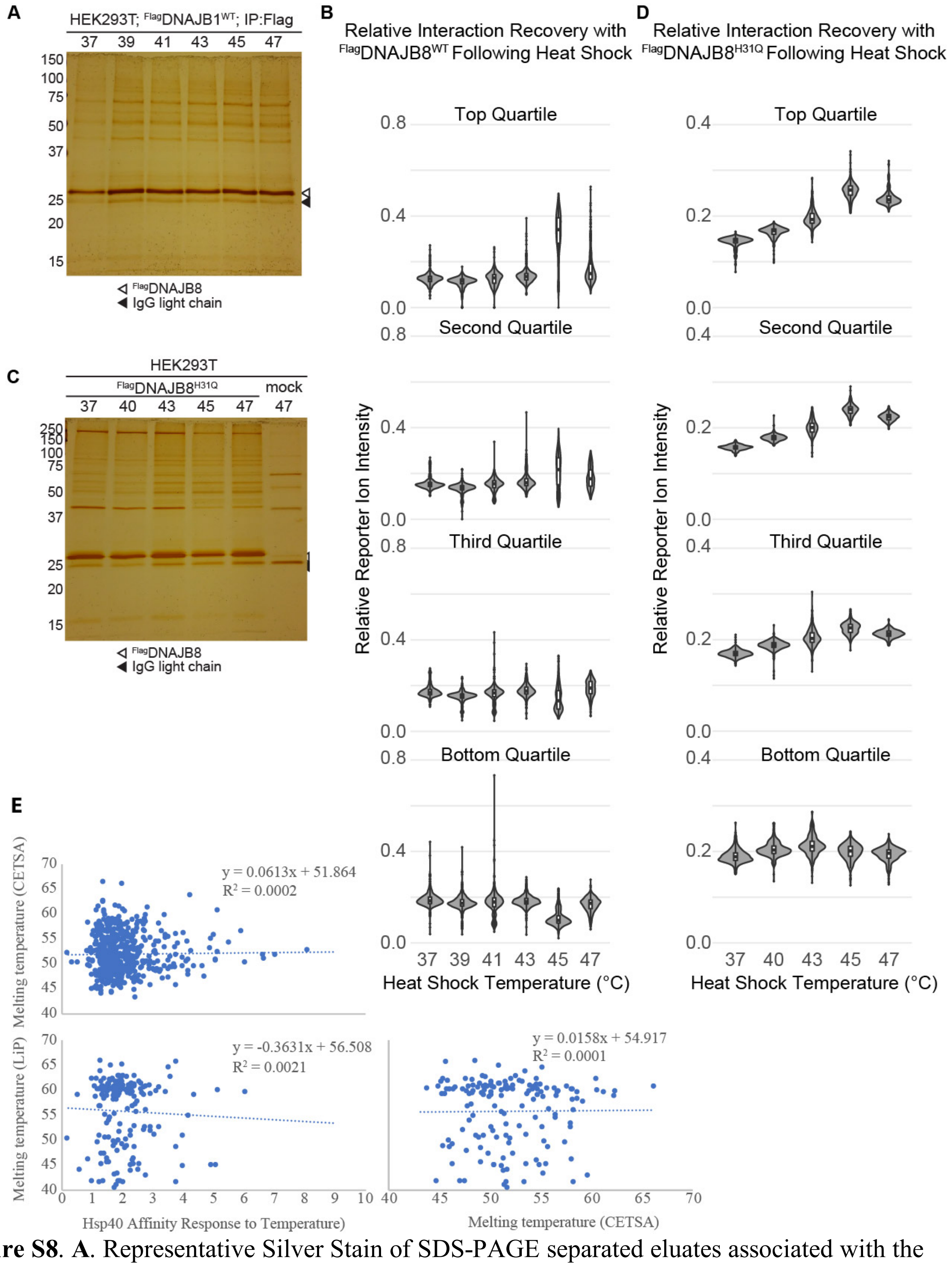


**Figure S8**. **A**. Representative Silver Stain of SDS-PAGE separated eluates associated with the experiment depicted in Figure 6A for the bait DNAJB8$^{WT}$. **B**. Variation in relative reporter ion intensities with temperature for DNAJB8$^{WT}$ associated proteins, organized by quartiles. **C**. Representative Silver Stain of SDS-PAGE separated eluates associated with the experiment depicted in Figure 6A with the bait being DNAJB8$^{H31Q}$. **D**. Variation in relative reporter ion intensities with temperature for DNAJB8 $^{H31Q}$ associated proteins, organized by quartiles. Note that due to the data covering a smaller range, different scale bars are used here than in Figure S8B. **E**. Relationship between increased Hsp40 affinity with heat shock, and melting temperature as determined by CETSA experiments in Jurkat cells[1] or limited proteolysis (LiP) in HeLa cells[2].
